# Environmental DNA metabarcoding provides enhanced detection of the European eel *Anguilla anguilla* and fish community structure in pumped river catchments

**DOI:** 10.1101/2020.07.22.216523

**Authors:** Nathan P. Griffiths, Jonathan D. Bolland, Rosalind M. Wright, Leona A. Murphy, Robert K. Donnelly, Hayley V. Watson, Bernd Hänfling

**Author notes:** **Correspondence**, Nathan P. Griffiths, Hardy Building, University of Hull, Hull, HU6 7RX, UK. **Funding information** Environment Agency, UK; Internal Drainage Boards, UK; University of Hull, UK.

## Abstract

The European eel *Anguilla anguilla* (eel hereafter) is critically endangered and has a catadromous lifecycle, which means adult eels that live in pumped catchments must pass through pumps during their downstream spawning migration. We are currently lacking detailed site-by-site eel distribution information in order to estimate the overall impact of individual pumping stations on eel escapement, and as such lack the data to enable informed prioritisation of pumping station management and targeted mitigation. In this study, we investigated whether environmental DNA (eDNA) metabarcoding can provide increased detection sensitivity for eel and fish community structure in highly regulated pumped catchments, when compared directly to current standard practice fish survey protocols (seine netting/electric fishing). Eels were detected in 14/17 sites (82.4%) using eDNA metabarcoding in contrast to 3/17 (17.6%) using traditional catch methods. Additionally, when using eDNA monitoring species richness was higher in 16/17 sites (94.1%) and site occupancy ≥ traditional methods for 23/26 of the fish species detected (88.5%). While eDNA methods presented significantly higher average species richness and species site occupancy overall, eDNA and Catch methods were positively correlated in terms of species richness and site occupancy. We therefore found that eDNA metabarcoding was a high sensitivity method for detecting eels in pumped catchments, while also increasing the detection of overall fish community structure compared to traditional catch methods. In addition, we highlight how eDNA monitoring is especially suited to increased detection of particular species, with traditional methods sufficient for others. This high sensitivity, coupled with the ability to sample multiple sites in a short time frame suggests eDNA metabarcoding could be an invaluable tool when prioritising pumping station management.

## 1 INTRODUCTION

The European eel (*Anguilla anguilla*) is a critically endangered catadromous fish species which has faced significant declines in recent decades (Bilotta *et al*., 2011; Jacoby & Gollock, 2014; Podgorniak *et al*., 2016; Correia *et al*., 2018). This marked decline has resulted in specific EU legislation, requiring member states to adopt eel management plans (The EC Eel Regulation (1100/2007)). These regulations aim to promote recovery by allowing >40% of the historic eel biomass prior to anthropogenic impacts passage from inland waters to the sea to facilitate spawning activity (Aalto *et al*., 2016). Despite these measures, The International Council for the Exploration of the Sea (ICES) Working Group on Eels (WGEEL) reports that current *A. anguilla* recruitment remains consistently <2% in recent years, with recruitment at 1.9% and 1.4% in 2018 and 2019 respectively (ICES, 2019). Such declines, at least in part, are a consequence of anthropogenic impacts on rivers - the focus here being societal reliance on land-drainage pumping stations for water level management (Solomon & Wright, 2012; Buysse *et al*., 2014, 2015; Bolland *et al*., 2019). These structures operate by pumping water against the natural gradient to a higher downstream river elevation, regulating river levels in the upstream catchment. This is a requirement in many areas across the world to enable flood management, agricultural water supply, and navigation (Solomon & Wright, 2012; Bolland *et al*., 2019; ICES, 2019). The overall ecological impacts of operating pumps are not fully understood however, and only recently have concerns regarding their adverse impacts on eels and whole fish communities been highlighted (Solomon & Wright, 2012; Buysse *et al*., 2014, 2015; Bolland *et al*., 2019).

The ecology of *A. anguilla* makes this species particularly vulnerable to adverse impacts at pumping stations. As a catadromous species, *A. anguilla* must undertake two transatlantic migrations between their European inland/estuarine occupancy range and spawning grounds located in the Sargasso sea (Bonhommeau *et al*., 2008; Podgorniak *et al*., 2016; Correia *et al*., 2018). This life cycle means that in pumped catchments, mature eels must pass through pumps in order to achieve escapement and spawn. It is the necessity to pass through pumps, in addition to their elongated morphology that makes *A. anguilla* especially susceptible to entrainment at pumping stations (Buysse *et al*., 2014; Bolland *et al*., 2019). Buysse *et al*. (2014) found that mortality rates were 97 ± 5% for a propeller pump, 17 ± 7% for a large Archimedes screw pump, and 19 ±11% for a small Archimedes screw pump respectively - indicating that mortality rates differ between pump types. However, Bolland *et al*. (2019) highlighted the importance of accounting for indirect impacts such as reduced fitness, delayed migration and increased predation when coming into contact with pumps, which may prevent successful spawning. ICES (2019) provided a first estimate of the total loss of eel biomass attributed to hydropower and land-drainage pumps, estimated at 444.4 tonnes per year in the UK alone, with a ‘minimum estimate’ of 1625.8 tonnes across Europe. While these figures suggest clear adverse impacts of pumping, given the increased likelihood of flood events predicted under future climate change scenarios (Team *et al*., 2014), we will likely be increasingly reliant on these pumps for land drainage in the coming years. To mitigate this, highly efficient, non-delayed and safe downstream eel passage routes must be provided for pumped catchments that contain *A. anguilla*.

Pumping stations regulate rivers where *A. anguilla* were undoubtedly once present but flood risk management infrastructure (flood banks / levees, pipework and pumps) can present a complete barrier to upstream migrating eels. We are currently lacking detailed site-by-site fish community information required to estimate the overall impact of individual structures on eel spawning escapement, and as such lack the data to enable informed prioritisation of pumping station management and perform targeted mitigation (ICES, 2019). Knowledge of the eel distribution and fish community present at these sites is therefore valuable to water managers (Solomon & Wright, 2012), yet due to sampling difficulties the probability of detecting rare and elusive species using traditional methods is low - particularly in large river systems (Pont *et al*., 2018). However, numerous studies in freshwater habitats have demonstrated that environmental DNA (eDNA) monitoring methods can achieve higher detection sensitivity than traditional monitoring techniques (Hänfling *et al*., 2016; Pont *et al*., 2018; Strickland & Roberts, 2018; Itakura *et al*., 2019; McDevitt *et al*., 2019). These molecular monitoring methods are ideal for detecting species with patchy distributions or low abundances, often overlooked by traditional catch methods (Turner *et al*., 2015). Recent developments mean that PCR-based metabarcoding of eDNA is now considered a powerful tool for monitoring entire ecological communities (Deiner *et al*. 2017; Hering *et al*. 2018), enabling vast amounts of data acquisition from a single sampling visit. In addition, it has been reported that eDNA is applicable to river systems, yielding higher detection rates, less sensitivity to sampling conditions and increased efficiency (Pont *et al*., 2018; Strickland & Roberts, 2018). This suggests eDNA could be a useful tool for screening species composition in pumped catchments – enabling multiple sites to be screened in a single survey.

In this study, we investigated whether eDNA metabarcoding (Hänfling *et al*., 2016; Bylemans *et al*., 2018; Pont *et al*., 2018; Li *et al*., 2019) can be used as a tool to monitor eel and fish presence in pumped catchments. While the application of species-specific qPCR methods may reduce the likelihood of false negatives by avoiding “species masking” effects (Harper *et al*. 2018), the holistic understanding that is provided by eDNA metabarcoding is integral to better inform management decisions (Ruppert *et al*. 2019). If successful, eDNA metabarcoding could be applied to enable evidence-based management of pumping stations, facilitating the attainment of policy-based objectives and conservation targets going forwards. However, it is important that this method is validated in such fragmented lotic systems with highly regulated catchments and flows. Therefore here, eDNA metabarcoding data are directly compared to data from standard practice traditional fish capture methods (seine netting/electrofishing) gathered from the same sites in the same year. We hypothesised that eDNA metabarcoding will enable increased detection sensitivity for our target species *A. anguilla*, while also increasing our coverage of whole fish communities (indicated by species richness and species site occupancy). Furthermore, we expected eDNA and Catch methods would be positively correlated for species richness and species site occupancy, indicating agreement between the methods, and thus continuity in regard to decision making.

## 2 MATERIALS AND METHODS

### 2.1 Study sites

This study was carried out in a low lying and heavily pumped section of the Fens, UK (Figure 1). Our study system ‘The Middle Level’ is a complex network of heavily pumped waterways and drainage ditches. Due to peat shrinkage caused by historic land drainage, ground levels have continued to sink meaning that there is no longer a possibility for gravity drainage at the 70+ pumping stations required to drain the area (Solomon and Wright 2012). Thus, all water transfer from this catchment to the sea is pumped, often passing through multiple pumping stations as it is moved through the system. There is little/no natural flow in the absence of pumping, meaning this system transitions between a lentic and lotic state temporally. The 17 study sites had a routine fish survey (performed by the Environment Agency between 07/04/2017 and 25/07/2017; seine netting = 15, electric fishing = 2 (Table 2, Supporting information)) and an eDNA survey (collected between 23/10/2017 and 28/11/2017; 5 water samples at each site (plus field blank)) carried out at identical points, allowing a direct comparison between methods that same year. The eDNA sampling was carried out later to enable the catchment to return to baseline conditions following the traditional survey season, and to ensure eDNA detection was not biased by an influx of glass eel recruitment which peaks between March and June in the region (Kroes *et al*. 2020).

**Figure 1.**
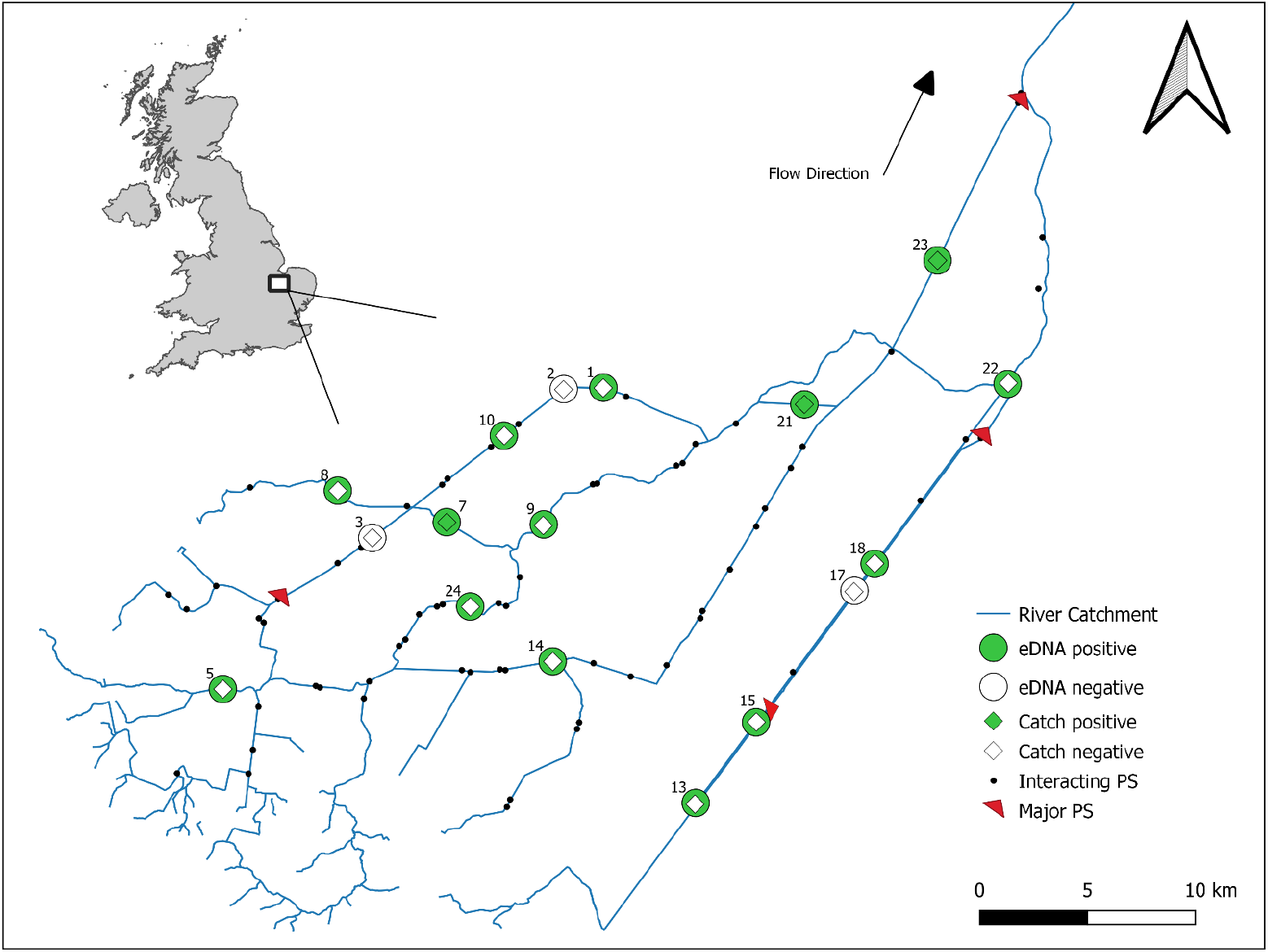
A map depicting the presence/absence and overall distribution of *Anguilla anguilla* across study sites based on eDNA surveys (circle) and traditional catch surveys (diamond). Net flow direction is indicated by the black arrow, while the pumping direction of major pumping stations (PS) is indicated by the direction of red triangles. Interacting pumping stations are also highlighted here (black dot), these pumps are connected to the main channel but pump drainage ditches into the river catchment (blue line), and thus are unlikely to entrain species present at our survey sites.

### 2.2 Water sampling

Five 2L surface water samples were taken at each site using sterile Gosselin™ HDPE plastic bottles (Fisher Scientific UK Ltd, UK). Each 2L sample consisted of 5 x 400ml sub-samples taken a few metres apart to account for the stochastic distribution of eDNA. Samples were taken by hand from a small inflatable boat, sterile gloves were worn by the sampler and changed between samples, the boat, oars, and waders were cleaned with bleach (10%), rinsed, then sprayed with Virkon (Antec International) between sites in order to prevent cross-site contamination. Samples were taken starting at the downstream end, then working upstream in 150m intervals with the mid-point being the national grid reference where traditional catch surveys were conducted. For each site, a 2L field blank (purified water) was included and handled alongside eDNA water samples to monitor for contamination.

Upon collection, water samples were stored on ice in a bleach-sterilised cool box during transit and taken back to our dedicated eDNA facility at the University of Hull for filtration. All samples and blanks were vacuum-filtered within 24 hours of collection. All surfaces and equipment were sterilised using 10% v/v chlorine-based commercial bleach solution (Elliott Hygiene Ltd, UK). Filtration equipment was immersed in 10% bleach for 10 minutes, soaked in 5% v/v MicroSol detergent (Anachem, UK) for an additional 10 minutes, then rinsed thoroughly with purified water between filtration runs. Whenever possible, the full 2L of water was vacuum-filtered through sterile 0.45μm cellulose nitrate membrane filters with pads (47mm diameter; Whatman, GE Healthcare, UK) using Nalgene filtration units - with 2 filters per sample to reduce filter clogging. Filters were then removed from units using sterile tweezers, placed in sterile 50mm petri dishes (Fisher Scientific UK Ltd, UK), sealed with parafilm (Sigma-Aldrich^®^, UK), and stored at −20°C until extraction.

### 2.3 DNA extraction

DNA was extracted from filters at the University of Hull dedicated eDNA facility in a designated sterile extraction area using the DNeasy PowerWater Kit (QIAGEN, Germany) following the manufacturer’s protocol. The duplicate filters from each sample were co-extracted by placing both filters back-to-back in a single tube for bead milling. Following extraction, the eluted DNA extracts (100μL) were quantified on a Qubit™ 3.0 fluorometer using a Qubit™ dsDNA HS Assay Kit (Invitrogen, UK) to confirm DNA was successfully isolated, then stored at −20°C.

### 2.4 eDNA metabarcoding

The eDNA library preparation and metabarcoding workflow applied here follows that outlined in Harper *et al*. (2019), but with the following modifications: the first PCR used 2μL of template DNA, 7μL of ddH20, and 0.5μL of BSA (Q5 2x High Fidelity Master Mix and primer volumes remained unchanged). The second PCR used 4μL of template DNA and 15μL of ddH20 (Q5 2x High Fidelity Master Mix and primer volumes remained unchanged). The second PCR thermocycling profile was also adapted as follows: 95°C for 3 mins, 10 cycles of 98°C for 20s and 72°C for 1 min, followed by a final elongation step at 72°C for 5 mins. The workflow is summarised below:

Nested metabarcoding using a two-step PCR protocol was performed, using multiplex identification (MID) tags in the first and second PCR step to enable sample identification as described in Kitson *et al*. (2019). The first PCR was performed in triplicate (i.e. 3× PCR replicates), amplifying a 106bp fragment using published 12S ribosomal RNA (rRNA) primers 12S-V5-F (5’-ACTGGGATTAGATACCCC-3’) and 12S-V5-R (5’-TAGAACAGGCTCCTCTAG-3’) (Kelly *et al*. 2014; Riaz *et al*. 2011). These selected primers have been previously validated, in silico, in vitro and in situ for UK freshwater fish species showing that all UK freshwater species can be detected reliably with the exceptions of distinctions between: *Lampetra planeri* / *Lampetra fluviatilis, Perca fluviatilis* / *Sander lucioperca*, three species of Asian carp (*Hypophthalmichthys nobilis, H. molitrix, Ctenopharyngodon idella*), and species within the genera *Salvelinus* and *Coregonus* (Hänfling et al. 2016). In our study, Lamprey were therefore assigned only to genus level, and Percidae assumed to be *P. fluviatilis*. PCR negative controls (MGW) were used throughout, as were positive controls using DNA (0.05ng/μL) from the non-native cichlid *Maylandia zebra*. The three replicates from the first PCR were pooled to create sub-libraries and purified with MagBIND^®^ RxnPure Plus magnetic beads (Omega Bio-tek Inc., GA, USA), following a double size selection protocol (Quail *et al*., 2009). Based on the ratios outlined in Harper *et al*. (2019), ratios of 0.9× and 0.15× magnetic beads to 100μL of amplified DNA from each sub-library were used. Following this, a second shuttle PCR was performed on the cleaned product to bind Illumina adapters to the sub-libraries. A second purification was then carried out on the PCR products with Mag-BIND^®^ RxnPure Plus magnetic beads (Omega Bio-tek Inc., GA, USA). Ratios of 0.7× and 0.15× magnetic beads to 50μL of each sub-library were used. Eluted DNA was then refrigerated at 4°C until quantification and normalisation. Once pooled, the final library was then purified again (following the same protocol as the second clean-up), quantified by qPCR using the NEBNext^®^ Library Quant Kit for Illumina^®^ (New England Biolabs^®^ Inc., MA, USA), and verified for fragment size (318bp) and purity using an Agilent 2200 TapeStation with High Sensitivity D1000 ScreenTape (Agilent Technologies, CA, USA). Once verified, the library was loaded (mixed with 10% PhiX) and sequenced on an Illumina MiSeq^®^ using a MiSeq Reagent Kit v3 (600-cycle) (Illumina, Inc., CA, USA) at the University of Hull. Raw sequence output was demultiplexed using a custom Python script, and our in-house bioinformatics pipeline metaBEAT v0.97.13 (https://github.com/HullUni-bioinformatics/metaBEAT) was used for quality trimming, merging, chimera removal, clustering, and taxonomic assignment of sequences against our curated UK fish reference database (Hänfling *et al*. 2016). Taxonomic assignment here used a lowest common ancestor approach based on BLAST matches that matched our reference database with minimum identity set at 98%.

### 2.5 Data analysis

During downstream analysis, data were analysed and visualised using R Version 3.6.3 (R Core Team, 2020). Reads assigned to family and genera containing only a single UK species were manually reassigned and merged with that species. In order to reduce the likelihood of eDNA false positives, blanks were used throughout and a low-read frequency threshold applied. All field/filtration blanks, PCR negative, and PCR positive controls were negative for *A. anguilla*, and the threshold applied at 0.001 to remove any reads making up less than 0.1% of total reads as previously applied with this 12S marker (Hänfling *et al*. 2016; Handley *et al*. 2019).

Samples of interest for this study (N=85) were then subset, and the mean number of reads for each site (N=17) calculated based on the five samples per site. Initially, data from traditional catch surveys were converted into ‘percentage catch’ for each species at each site. This enabled direct visual comparisons of relative abundance between methods, based on ‘percentage reads’ and ‘percentage catch’, visualised as bubble plots using ggplot2 3.2.0 (Wickham, 2016). Differences between survey methods were then compared statistically, based on the species richness and species occupancy obtained. Richness and Occupancy data were screened for normality, in order to meet assumptions a paired t-test was applied to species richness data, and a paired Wilcoxon signed rank test to species site occupancy data to test for significant differences between survey methods (McDonald, 2014). Correlations between eDNA and traditional catch methods were then tested for using Pearson’s and Spearman’s tests, for richness and site occupancy respectively (McDonald, 2014), and visualised using ggpubr (Kassambara, 2020).

## 3 RESULTS

### 3.1 Eel distribution

The total average eDNA reads across all study sites was 842,159, of which 8,830 (1.05%) were assigned to *A. anguilla*. The total number of individual fish caught using traditional catch methods across study sites was 14,102, of which 3 (0.02%) were *A. anguilla*. This comparison of survey methods shows that eDNA metabarcoding yielded an overall higher detection rate for *A. anguilla* than traditional catch methods. The eDNA metabarcoding approach detected *A. anguilla* in 14/17 (82.4%) of the sites surveyed, whereas traditional methods captured *A. anguilla* in 3/17 (17.6%) of sites (Figure 1). Both approaches tested positive for *A. anguilla* in three sites (7, 21 and 23), and both negative for another three sites (2, 3, and 17); agreement between methods in 6/17 sites (35.3%). The other 11/17 (64.7%) sites however did not agree, with traditional methods capturing no *A. anguilla* in sites where eDNA surveys tested positive.

### 3.2 Species composition

Overall, 26 fish species were detected in this study; eDNA metabarcoding detected 25/26 fish species (96.2%) across the 17 study sites, whereas traditional methods captured 16/26 fish species (61.5%). Of the 26 species detected, 15 (57.7%) were detected in both eDNA and traditional catch methods, whereas 10 (38.5%) were detected only using eDNA and one (3.85%) species was detected only using traditional capture methods (Figure 2). There were visual consistencies between species with higher % Reads (eDNA) and % Catch (traditional methods), including *Abramis brama, Esox lucius, Perca fluviatilis* and *Rutilus rutilus* (Figure 2).

**Figure 2.**
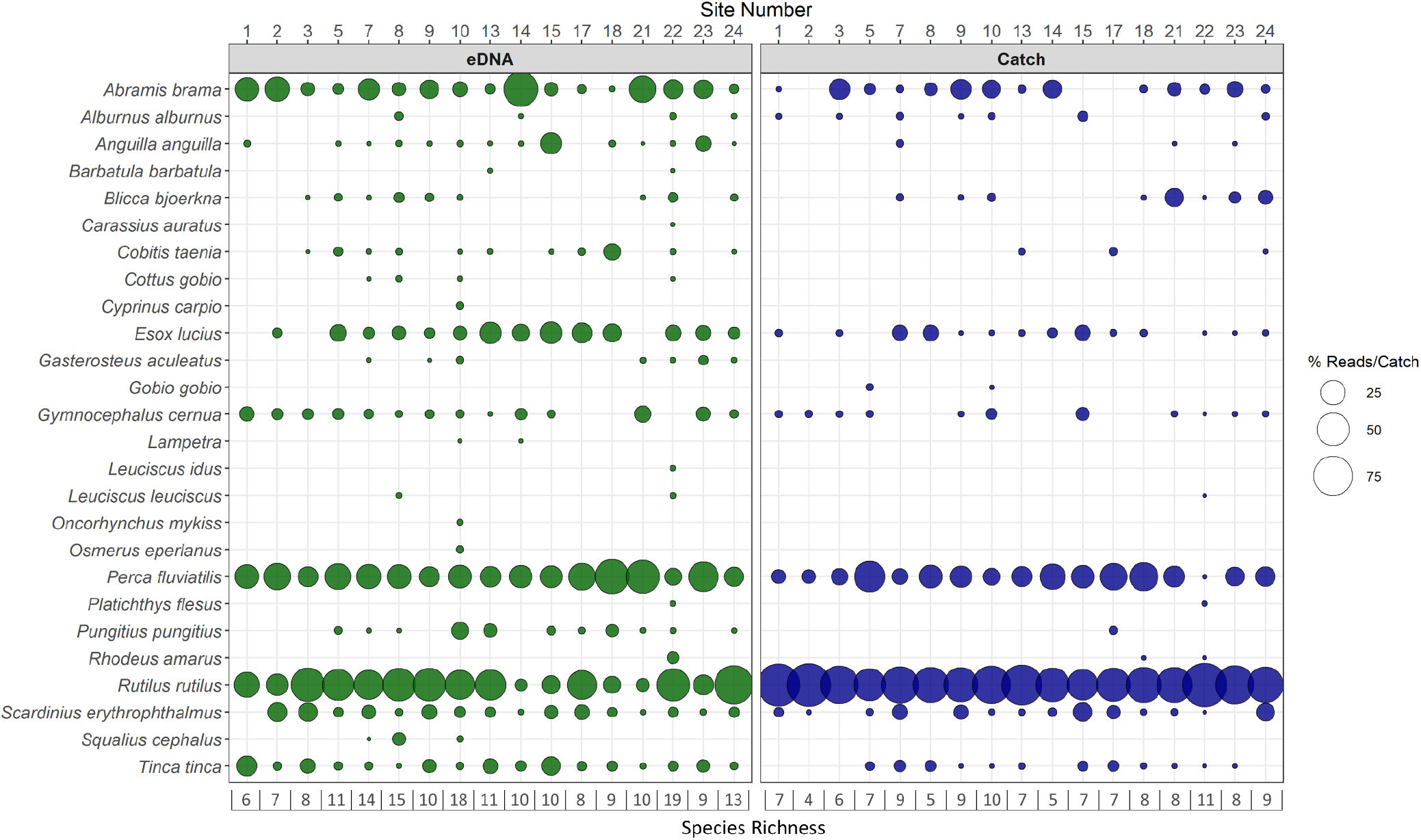
Bubble plot depicting % Reads/Catch (circle size) of each species detected at each site using eDNA metabarcoding (green fill, left) and traditional catch (blue fill, right). Presenting a visual comparison of relative abundance between methods, with total species richness at each site on the bottom row.

### 3.3 Species richness

eDNA metabarcoding yielded a higher species richness than traditional catch methods in 16/17 sites, with the exception being site 1. Additionally, eDNA (M=11.06) reported significantly higher mean species richness than traditional methods (M=7.47) (paired t-test; t = −5.0355, df = 16, p = 0.0001) (Figure 3a). While a Pearson’s product-moment correlation test showed the two methods were significantly correlated (R = 0.6, p = 0.011) (Figure 3b).

**Figure 3.**
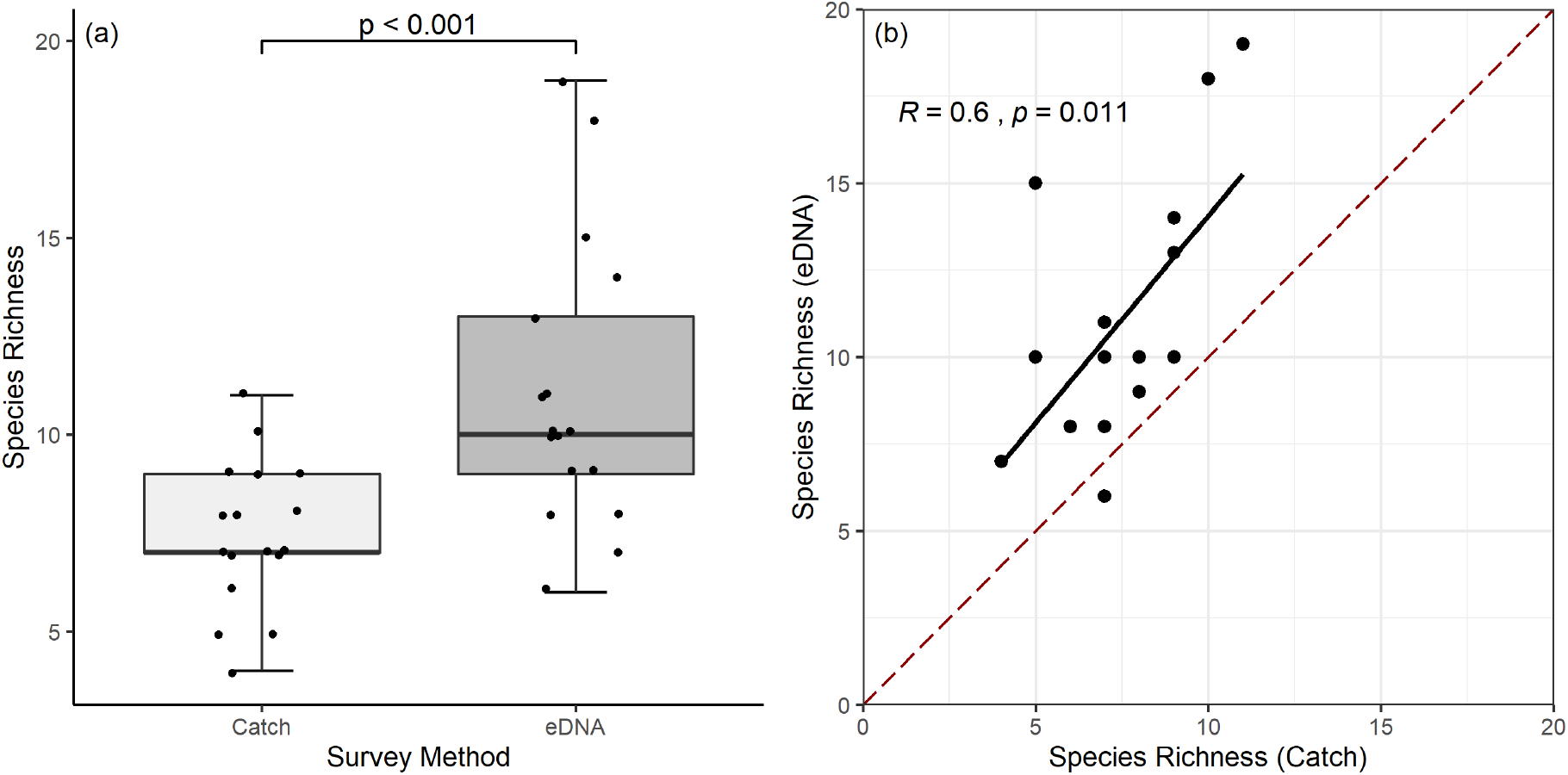
Box plot (a) of species richness at each site for each survey method, including p-value (paired t-test). Scatter plot (b) comparing site species richness between eDNA and traditional methods, including the Pearson correlation test output and associated regression line to visualise correlations between methods. The dashed red line indicates species richness equilibrium, where points above the line had a higher species richness using eDNA, while points below had higher Catch species richness.

### 3.4 Species site occupancy

eDNA metabarcoding reported a site occupancy ≥ traditional catch methods for 23/26 of the fish species detected, with *Alburnus alburnus, Rhodeus amarus* and *Gobio gobio* being the exceptions (Table 1). Additionally, eDNA (M=7.23) reported significantly higher mean site occupancy than traditional methods (M=4.88) (paired Wilcoxon test; V = 29.5, p = 0. 001607) (Figure 4a). While a Spearman’s rank correlation test showed the two methods were significantly correlated (R = 0.76, p = 5.762e-06) (Figure 4b).

**Figure 4.**
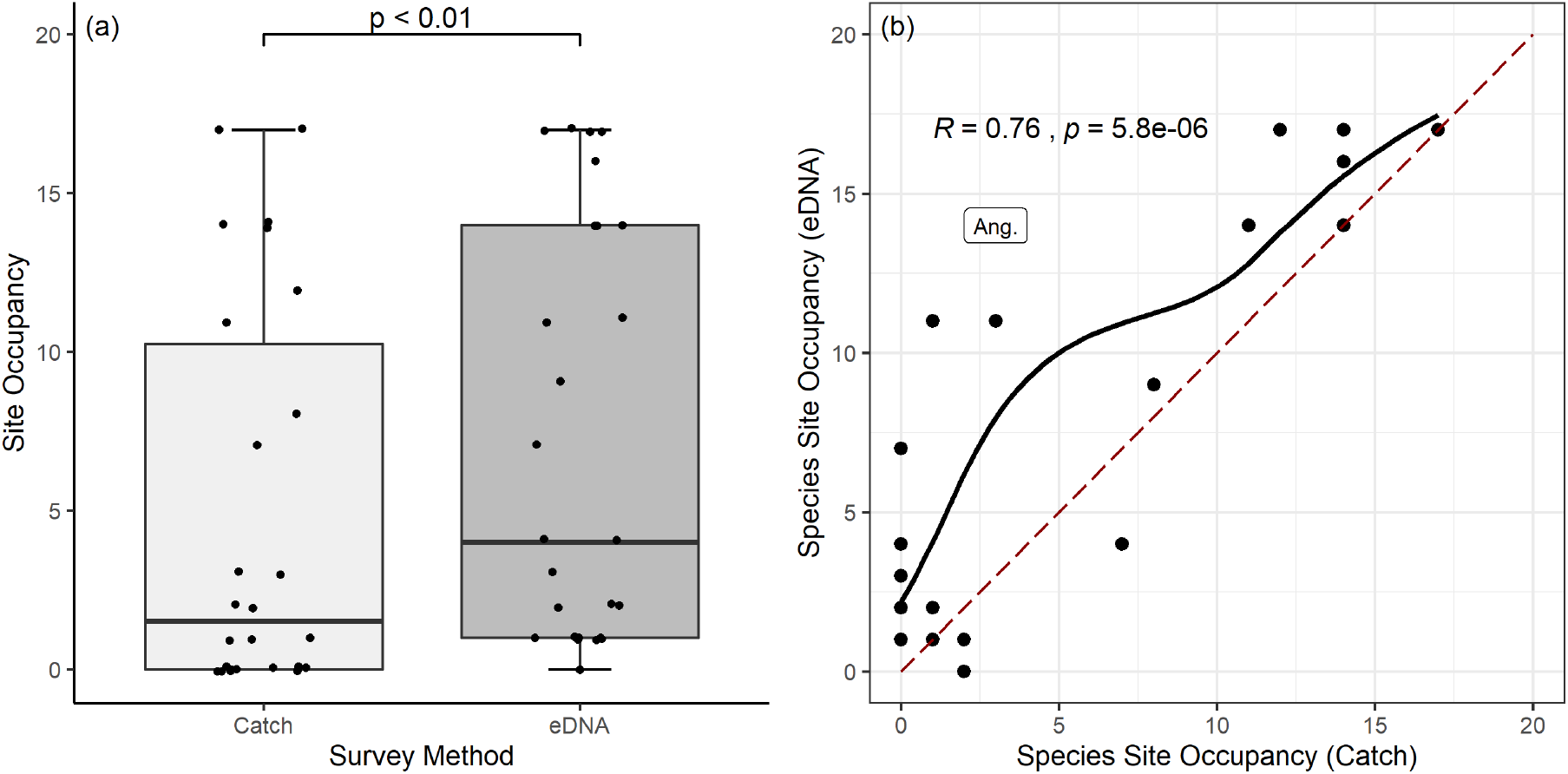
Box plot (a) of species site occupancy at each site for each survey method, including p-value (paired Wilcoxon test). Scatter plot (b) comparing species-specific site occupancy between eDNA and traditional methods, including the Spearman’s rank correlation test output, and a smooth curve (loess) to visualise associations. The dashed red line indicates site occupancy equilibrium, where points above the line indicate a species was detected at more sites using eDNA and points below the line indicate a species was captured at more sites using traditional methods (text label indicates our target species *A. anguilla*).

**Table 1.** A comparison of the species site occupancy obtained from eDNA metabarcoding and traditional catch methods. In the species column, UK BAP (Biodiversity action plan) species (+) and non-native/introduced species (*) are indicated.

| **Species** | **eDNA** | **Catch** |
| --- | --- | --- |
| *Abramis brama* | 17 | 14 |
| *Alburnus alburnus* | 4 | 7 |
| *Anguilla anguilla* + | 14 | 3 |
| *Barbatula barbatula* | 2 | 0 |
| *Blicca bjoerkna* | 9 | 8 |
| *Carassius auratus* * | 1 | 0 |
| *Cobitis taenia* + | 11 | 3 |
| *Cottus gobio* | 4 | 0 |
| *Cyprinus carpio* * | 1 | 0 |
| *Esox lucius* | 14 | 14 |
| *Gasterosteus aculeatus* | 7 | 0 |
| *Gobio gobio* | 0 | 2 |
| *Gymnocephalus cernua* | 14 | 11 |
| *Lampetra* + | 2 | 0 |
| *Leuciscus idus* * | 1 | 0 |
| *Leuciscus leuciscus* | 2 | 1 |
| *Oncorhynchus mykiss* * | 1 | 0 |
| *Osmerus eperlanus* + | 1 | 0 |
| *Perca fluviatilis* | 17 | 17 |
| *Platichthys flesus* | 1 | 1 |
| *Pungitius pungitius* | 11 | 1 |
| *Rhodeus amarus* * | 1 | 2 |
| *Rutilus rutilus* | 17 | 17 |
| *Scardinius erythrophthalmus* | 16 | 14 |
| *Squalius cephalus* | 3 | 0 |
| *Tinca tinca* | 17 | 12 |

## 4 DISCUSSION

To our knowledge, this study is the first to validate eDNA metabarcoding of fish communities specifically within heavily regulated pumped river catchments. We found that eDNA metabarcoding consistently outperformed standard practice methods, in terms of increased detection for our target species *A. anguilla* and enhancing fish community structure knowledge at our study sites, while revealing presence of additional elusive species. Here, we discuss these findings and what they mean for eDNA as a tool for stakeholders to inform management decisions in pumped river catchments.

### 4.1 Detecting *A. anguilla* in pumped catchments

When considering pumping station management *A. anguilla* is a key species given they are critically endangered and have specific legislation to protect them from human-mediated activities (Council Regulation (EC) No. 1100/2007) (Buysse *et al*., 2014, 2015; Bolland *et al*., 2019). In this study, all three sites where traditional methods captured *A. anguilla* had positive eDNA signals, while the three eDNA negative sites were also negative for traditional catch methods. Most notably, eDNA metabarcoding detected *A. anguilla* at 11 additional sites where traditional capture methods did not. We therefore conclude that eDNA metabarcoding was more sensitive for detecting *A. anguilla* in managed pumped catchments than traditional methods. This was not unexpected, given the documented challenges in sampling eels from aquatic environments using seine netting and electric fishing (Naismith & Knights, 1990; Degerman *et al*., 2019). Similarly, a recent study by Itakura *et al*. (2019) found that single species qPCR based eDNA monitoring had a greater detection sensitivity for the Japanese eel *Anguilla japonica* than electrofishing in rivers in Japan. One consideration in our study however, is that silver eel migration generally begins in autumn when water temperature decreases (Acou *et al*. 2008), this corresponds with our eDNA sampling and could potentially enhance detection as these mature eels migrate downstream. Based on our results, we recommend eDNA metabarcoding as a complimentary/alternative method to traditional catch when eel presence/absence data is required for informing management decisions in pumped catchments.

### 4.2 Fish community detection in pumped catchments

While the priority species for this study was *A. anguilla*, the composition and structure of the fish community as a whole remains an important factor in management decision making and is of interest to wider stakeholders (Solomon & Wright, 2012; Pont *et al*., 2019). It is therefore important that eDNA monitoring is able to produce data on these should it be implemented as a standardised monitoring framework.

We observed that eDNA methods were able to detect significantly more species in pumped catchments than standard practice catch methods, as indicated by the total number of species detected (25 and 16, respectively) and increased species richness at individual sites. Previous studies have clearly shown that eDNA metabarcoding is a highly sensitive method for detection of freshwater fish which outperforms traditional survey techniques in lentic environments (Hänfling *et al*., 2016; Handley *et al*., 2019; Li *et al*., 2019). This study adds to the mounting evidence that this is also true in lotic environments with unregulated (Bylemans *et al*., 2018; Pont *et al*., 2018) and regulated flows (McDevitt *et al*., 2019). Despite the underestimation of true species richness in traditional surveys, when compared to eDNA metabarcoding there was a strong positive correlation between both methods, and thus the relative importance of sites based on species richness for both methods were related.

Of course, not all species are equally weighted when it comes to making management decisions (Solomon & Wright, 2012; Nunn *et al*., 2014; Beng & Corlett, 2020; Sepulveda *et al*., 2020), and so it is important to consider the potential for any method biases (preferential detections or underrepresented species). We found that overall eDNA had significantly higher mean site occupancy than traditional methods, and thus was more sensitive to individual species detections, while site occupancy was positively correlated between methods. Here we observed that species missed by catch methods were low in percentage reads and site occupancy using eDNA methods (Figure 2), suggesting low abundance species may be overlooked by catch methods, whereas agreement between methods is higher for abundant species (Figure 4b). This increased sensitivity could enable a more targeted focus to conservation species management, enabling increased detection of priority species such as *Cobitis Taenia* notable in our study (Nunn *et al*., 2014). However, while eDNA site occupancy was ≥ traditional methods for 23/26 of the fish species detected. Two species *A. alburnus* and *R. amarus* had a higher detection rate using traditional methods, while *G. gobio* was not detected using eDNA metabarcoding at all, despite being identified on two occasions using traditional methods. While this could be due to the morphological identification bias of traditional surveys (Li *et al*., 2019), the influences of reference databases and species ecology should be considered with eDNA metabarcoding in such instances (Bylemans *et al*., 2018). In our case primers and reference databases had been previously validated showing that this species can be reliably detected (Hänfling *et al*. 2016). Furthermore, while *G.gobio* was not detected in this study, it was present in other samples in this workflow which were not part of this study. This suggests species ecology, sampling conditions, or misidentification from catch surveys as possible explanations. Previous comparisons of detection rates from eDNA and traditional surveys have found that species detectability increases for both methods based on the density of target organisms (Hering *et al*. 2018). However, it must be considered that detectability remains imperfect for both methods. While the detection rate in our study was on average higher for eDNA this is not true for all species. We observed increased stochasticity for rare species with lower detection rates across methods, meaning that some species may genuinely have higher detection rates with conventional survey designs, or may be overlooked by both methods. It has been noted in the literature for example, that traditional surveys are prone to over-estimation of sub-surface species, and under-estimation of benthic and rare species when compared to eDNA (Pont *et al*. 2019). This could explain why eDNA was more sensitive for *A.anguilla* in our study, while traditional catch methods yielded higher detection for *A.alburnus*. Here, we highlight the importance of understanding potential limitations or biases of species detectability, prior to using any type of monitoring data to inform management decisions.

In order to enable direct comparisons between methods we used naïve occupancy, which does not account for imperfect detection (Ficetola *et al*. 2015). This can lead to underestimation of species distribution if replication is low (Sutter and Kinziger 2019), yet risk false positives if replication is large and false positive rate is high. While a conservative approach to occupancy can reduce false positives, many true positives are also discarded (Ficetola *et al*. 2015). Our study therefore applied a low-frequency threshold to reduce eDNA false positive rates, rather than use a conservative approach to occupancy and risk overlooking rare and elusive species. Based on this and our blanks, the potential for unfiltered false positives in our data is reduced. However, false positives due to environmental contamination remain a potential source of error which could be reflected in species detected in single sites. In our study these included *Carassius auratus, Leuciscus idus*, and *Oncorhynchus mykiss;* the species with only a single positive eDNA sample. These species are all considered non-native but introduced to the UK (Table 1), but this does not mean we can rule out environmental contamination, and as such decision-support schematics should account for such scenarios based on local management plans (Sepulveda *et al*. 2020).

### 4.3 Conclusions

While eDNA has been well validated in lentic systems (Hänfling *et al*., 2016; Handley *et al*., 2019; Li *et al*., 2019), it is acknowledged that downstream transportation of eDNA and fluctuations in river flow can influence eDNA detection and spatial resolution in lotic systems (Turner *et al*., 2015; Bylemans *et al*., 2018; Pont *et al*., 2018; Itakura *et al*., 2019; Milhau *et al*., 2019; Laporte *et al*., 2020). Pumped catchments are prone to binary fluctuations in flow, changing from a lentic to a lotic water body with pump operation to regulate river level (Solomon & Wright, 2012; Buysse *et al*., 2014; Bolland *et al*., 2019). As pump operation, in most instances, is influenced by rainfall, there are likely seasonal trends to consider as well as smaller scale temporal variation in pumping regimes with currently unknown influences on species detectability. It may be that pump operation acts as a conveyor belt for eDNA diversity, as described in (Deiner *et al*., 2016), and sampling at a pumping station after a pumping event can reduce eDNA stochasticity and reduce the number of spatial replicates required. Alternatively, small pumped catchments may be diluted by heavy rainfall events and pump operation flushing the system, reducing the detectability of prevailing fish. A study by Shogren *et al*. (2017) however, investigated the impact of eDNA transport in controlled streams - highlighting the complexities of applying predictive models to variable environments. In respect to our highly variable study site, any eel detected upstream of a sampling point would eventually have to pass through downstream pumps in order to achieve seaward migration, and so such effects would be negligible on the interpretation of results from a management perspective. These factors should however be considered in regard to study/sampling design, and complexity of the study system when making additional inferences from results in pumped catchments. As such, future research into the influence of seasonal and daily variation in pump activity on eDNA performance in pumped catchments is recommended.

While further consideration may be required before applying this method as a standardised tool in management frameworks, this study demonstrates successful application of eDNA as a high sensitivity tool to screen for eel and fish in pumped catchments. In addition to increased detection sensitivity, significant correlations between eDNA and Catch methods further evidence increased confidence in eDNA based decision making (Jerde, 2019). Validating eDNA metabarcoding as a non-invasive method with decreased sampling effort and higher detection rates of target species. Sepulveda *et al*. (2020) suggests that the reliability of eDNA methods is not a barrier, the problem is that we often lack the tools to integrate inherent uncertainty into decision-making frameworks. These molecular methods could be applied methodically to work with such frameworks, but should be integrated in a manner which could fill existing knowledge gaps while accounting for related uncertainty. We therefore conclude that this workflow could be optimised as a method to inform management for pumped catchments in Europe and beyond.

## Acknowledgements

We would like to thank landowners and managers for granting permission to access sampling sites, and the UK Environment Agency who shared their monitoring data with us. In addition, we are grateful to N. Baker and R. Ainsworth for assistance with fieldwork, and L. R. Harper, C. Di Muri, G. Sellers and M. Benucci for their support with bioinformatics and the handling of outputs.

## Contributions

N.P.G carried out bioinformatics, data analysis and manuscript preparation. L.A.M coordinated fieldwork and sample collection. R.K.D and H.V.W carried out lab work. J.B, R.W, and B.H conceived the study, acquired funding and supported with manuscript preparation/ideas.

## Data accessibility

Raw sequence reads have been archived on the NCBI Sequence Read Archive (SRA) under BioProject: PRJNA646357; BioSamples: SAMN15541168 - SAMN15541275; SRA accessions: SRR12232432 - SRR12232539. R scripts, Jupyter notebooks and corresponding data have been made available in a dedicated GitHub repository, which is permanently archived at (https://doi.org/10.5281/zenodo.3951418).

## Supporting Information

**Table 2.** An overview of Environment Agency Catch methods applied at each site, including Date, Area fished, Survey method, Survey strategy and Number of runs. This is tabulated and available as a .csv file.

